# Temperature-Switchable Genome Editors from Extremophile-Derived Integrases

**DOI:** 10.64898/2026.09.10.749218

**Authors:** Jack H. Arnold, Isabella G. Romano, Devyn L. Del Curto, Elise K. Wilbourn, Ellis L. Torrance, Valerie A. Fierro, J. Bryce Ricken, Kelly P. Williams, Joseph S. Schoeniger, Jesse L. Cahill

## Abstract

Integrases are site-specific recombinases encoded by phages and other mobile genetic elements. They mediate DNA integration, excision, and inversion between cognate attachment (*att*) sites. Although integrases are powerful tools for genetic engineering and synthetic biology, most systems lack intrinsic mechanisms that limit activity after expression, creating potential for unintended recombination. We hypothesized that extremophiles could provide temperature-responsive integrases because their enzymes evolved under selective pressure to operate within the thermal ranges experienced by their hosts. To test this concept, we linked integrase-*att* pairs from 458,683 prokaryotic genome assemblies to curated host growth-temperature metadata. This analysis revealed temperature-associated structure among integrase clusters and established a candidate pool for testing temperature-responsive recombinases. GC content in tyrosine integrase-associated *attB* sites showed a modest increase in higher-temperature hosts. We developed an inversion assay using a single-copy reporter plasmid and *sacB* counterselection to quantify integrase activity across temperatures. Thermophile-derived integrases from *Thermus thermophilus* and *Geobacillus stearothermophilus* displayed hot-ON/cold-OFF activity profiles, whereas an integrase derived from the psychrotroph *Pseudomonas cerasi* showed cold-ON/hot-OFF activity. Together, these results establish host thermal niche as a guide for discovering intrinsically temperature-switchable integrases and provide a foundation for engineering thermally controlled genome editing systems.

**GRAPHICAL ABSTRACT:** Discovery and validation of temperature-switchable integrases. **(A)** Integrase–attachment site pairs were identified from the TIGER/Islander resource and linked to host thermal metadata from curated growth-temperature databases. Candidate systems were grouped by host thermal class, including psychrophile, psychrotroph, mesophile, thermophile, and hyperthermophile sources. **(B)** Selected integrase-*att* pairs were tested in an *E. coli* inversion assay in which recombination switches a reporter cassette from a *sacB*-sensitive state to a sucrose-resistant state. **(C)** Thermophile-derived integrases exhibited hot-ON/cold-OFF recombination profiles, whereas a lower-temperature-associated *Pseudomonas cerasi* integrase showed cold-ON/hot-OFF activity, demonstrating that host thermal niche can guide discovery of temperature-responsive recombinases.

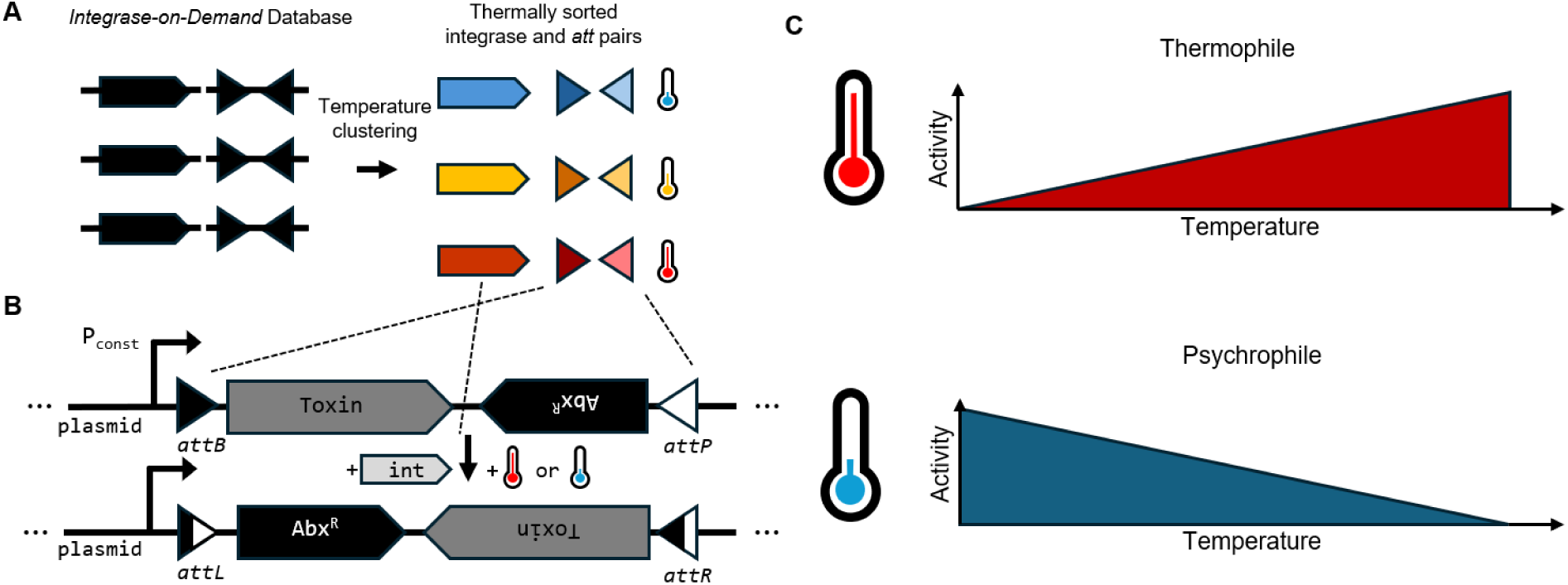

## INTRODUCTION

Genome editing technologies, including CRISPR-Cas systems and site-specific recombinases, have transformed genetic engineering across medicine, agriculture, biotechnology, and environmental science (1–5). However, as these tools are deployed in increasingly complex biological contexts, precise control over editor activity has become a central challenge. Prolonged or uncontrolled genome editor activity can be toxic, increase off-target modification, promote genomic instability, and produce unintended biological consequences (6–8). These properties also can make it challenging to assemble or deliver synthetic constructs that contain both genome editors and their targets, limiting digital synthetic biology opportunities. Existing control strategies, including inducible promoters, ligand-responsive systems, light-inducible proteins, and allosteric regulation, have enabled important advances but can be limited by leakiness, delivery constraints, incomplete suppression of background activity, or poor penetration in biological tissues (4,9,10). These limitations motivate the development of genome engineering tools with intrinsic mechanisms that restrict activity to defined conditions.

Temperature is an attractive control parameter for intrinsic biological activity because it is easily tunable, readily measured, and already used broadly to regulate biochemical activity. Temperature-dependent enzyme behavior is widespread in nature: thermophilic enzymes often maintain stability and catalytic function at elevated temperatures, whereas psychrophilic enzymes are adapted for activity at lower temperatures and frequently lose function as temperature increases (11,12). These properties have been exploited in biotechnology, most notably through thermostable DNA polymerases that enabled polymerase chain reaction and engineered polymerase variants with altered temperature sensitivity (13,14). More broadly, directed evolution and mutagenesis have generated biomolecules with altered thermal response profiles, including recombinases and transcriptional regulators with temperature-dependent activity (15,16). Together, these examples suggest that enzymes from organisms adapted to distinct thermal niches may provide useful starting points for developing temperature-responsive biological control systems.

Phage and mobile-element integrases are particularly compelling candidates for temperature-switchable genome engineering. Integrases are site-specific recombinases that catalyze recombination between defined DNA attachment (*att*) sites. In the canonical integration reaction, an integrase recombines a phage-associated *attP* site with a bacterial or chromosomal *attB* site, generating hybrid *attL* and *attR* sites that flank the integrated DNA (1,17). In many systems, the reverse reaction can be promoted by a recombination directionality factor or excisionase, enabling excision and regeneration of *attP* and *attB*. The same site-specific chemistry can also be adapted for genetic control: when compatible attachment sites are placed on the same DNA molecule, their orientation and position determine whether recombination results in integration, excision, or inversion of the intervening sequence. Thus, integrases are widely used in synthetic biology for genomic integration, gene regulation, and construction of genetic memory systems (18–20).

Unlike nuclease-based editing systems, integrase-mediated recombination does not require double-strand DNA open breaks, and many integrase reactions are intrinsically reversible (1). Despite these advantages, integrase activity is usually regulated indirectly through transcriptional control of integrase expression rather than through intrinsic control of enzyme activity. Productive recombination requires assembly of a multimeric synaptic complex, and even low levels of leaky expression may generate sufficient integrase activity to recombine available *att* sites (18,21). In addition, transcriptional shutoff does not immediately inactivate integrase protein that has already accumulated (22,23). As a result, expression-based control may be insufficient for applications in which integrases and their cognate *att* sites must coexist in a cell but remain inactive until a defined trigger is applied and then return to an inactive state to prevent additional recombination.

Extremophile-derived integrases provide an opportunity to address this limitation and utilize temperature as an intrinsic mechanism of control. Because integrases function within the cellular environments of their hosts, enzymes from thermophiles, psychrophiles, psychrotrophs, and related organisms may carry activity profiles shaped by host-specific thermal conditions (11,12). We therefore hypothesized that organisms adapted to distinct thermal niches could provide temperature-responsive integrases. Testing this hypothesis requires more than identifying integrase homologs from organisms with different growth temperatures. Integrase function depends on cognate attachment sites, so candidate discovery must preserve the relationship between each integrase and its predicted *att* sites (1) because there is a potential for *att* site properties such as GC content to influence the temperature dependence of activity. The TIGER/Islander (24) resource, which precisely maps genomic islands and associates integrase genes with their cognate attachment sequences, enables moving from sequence mining to experimentally testable integrase-*att* systems.

Here, we develop a framework that connects prokaryotic genome assemblies, host growth-temperature metadata, and precisely mapped integrase-attachment site pairs. This framework enables integrases to be analyzed in the context of host thermal niche, integrase family, sequence similarity, and cognate DNA target sites. More broadly, linking curated growth-temperature data to genome-scale resources potentially creates a foundation for identifying temperature-associated features across many biological components, not only integrases. We use this framework to select representative integrase-*att* pairs for experimental validation in a temperature-resolved recombination assay, establishing a route to discover genetic control elements whose activity is regulated by temperature rather than expression alone.

## MATERIALS AND METHODS

### Database construction and temperature annotation

To construct a comprehensive dataset of candidate integrases, we aggregated sequences from multiple publicly available and internal resources. The primary TIGER/Islander dataset (24) contains 1,757,053 precisely mapped genomic islands found among 459,935 bacterial and archaeal genome assemblies, with their associated *att* site and integrase sequences (448802 unique tyrosine integrase sequences, and 75588 unique large serine integrase sequences).

Organism-level temperature data were taken from several curated databases, including those from the Sauer (25) (11,104 entries) and Gosha (26) (14,433 entries) laboratories, TEMPURA (27) (8,639 entries), and the Joint Genome Institute Genomes OnLine Database (28) (JGI GOLD; 17,984 entries). Databases were selected based on their inclusion of experimentally determined or curated growth temperature parameters, including optimal growth temperature (OGT, °C), maximum growth temperature (T_max, °C), minimum growth temperature (T_min, °C), or categorical thermal classifications. Where possible, priority was given to numerical optimum growth temperatures over thermal classifications. To associate integrase sequences with host organism temperature profiles, records were merged using a hierarchical matching strategy. When available, the NCBI nine-digit genome assembly ID was used as the primary key for database integration. In cases where such identifiers were absent, species-level matching was performed using standardized taxonomy names. If neither an NCBI ID nor species-level taxonomic assignment was available, the data was not included in our final database.

Following integration of organism OGT, each integrase entry was annotated with temperature metadata corresponding to its host organism, enabling classification into the 5-category system as defined by the JGI GOLD database, where psychrophiles are those with OGT below 9.99°C, psychrotrophic organisms grow between 10 to 19.99°C, mesophiles grow between 20 and 45°C, thermophiles grow between 45.1 and 79.99°C, and hyperthermophiles grow above 80°C. Where multiple OGTs were reported, a median OGT was calculated with all numerical OGT values from all databases. This unified dataset provided the foundation for downstream analysis of temperature-dependent integrase activity and candidate selection for experimental validation.

Following database integration, all temperature metadata sources were combined into a unified dataset, and a comprehensive list of species was compiled alongside their associated genome assembly identifiers. Species entries were standardized and annotated using temperature information from all available sources, prioritizing records that contained at least one experimentally determined or curated optimal growth temperature. Organisms were subsequently classified into thermal categories according to the JGI classification framework, enabling consistent grouping into thermophilic, mesophilic, or psychrophilic classes. To ensure dataset integrity, duplicate species entries arising from overlap between databases were systematically removed using species-level de-duplication. This process resulted in a final curated dataset comprising 12,760 genome assemblies with associated temperature annotations, providing a robust foundation for downstream mapping of integrase sequences to host thermal profiles.

### GC content analysis

To further investigate sequence-level adaptations associated with thermal environments, we analyzed the GC content of *attB* attachment sites corresponding to integrases across classes range of optimal growth temperature annotations. Sequences for *attB* sites were taken from *Integrase-on-Demand* (29). Serine and tyrosine integrases were analyzed separately. Additionally, sites for serine recombinases with only the catalytic domain (typically resolvases or invertases) were excluded from analysis.

Linear relationships between host growth temperature and *attB* or genomic GC content were evaluated separately for serine and tyrosine integrase-associated *attB* sites and with or without mesophile class subsets (**Supp. Fig S1A-L**). Mesophile exclusion was used to avoid potential bias from the use of 37°C in experimental culture conditions where this temperature may not truly correspond to the OGT of an organism (**Supp. Fig S1M**). Host growth temperature was represented using the available numerical or categorical temperature annotation for each organism, prioritizing curated or experimentally reported OGT values where available. Pearson correlation coefficients (*r*) and statistical significance for *attB* or genomic GC and OGT were assessed using an ordinary least squares linear regression with a significance threshold of P<0.05, and results are reported as significant or not significant in **Table 1 and Supp. Table S1**. Statistical analyses for Pearson correlation coefficients were performed in Python using the Scipy.stats package version 1.18.0 (30). A Steiger’s Z-test (31) was performed to compare the dependent Pearson coefficients between *attB*-OGT (*r_attB_*_,OGT_), genomic GC-OGT (*r*_genome,OGT_), and *attB* GC-genomic GC (*r_attB_*_,genome_) using an online calculator (https://www.psychometrica.de/correlation.html).

**Table 1:**
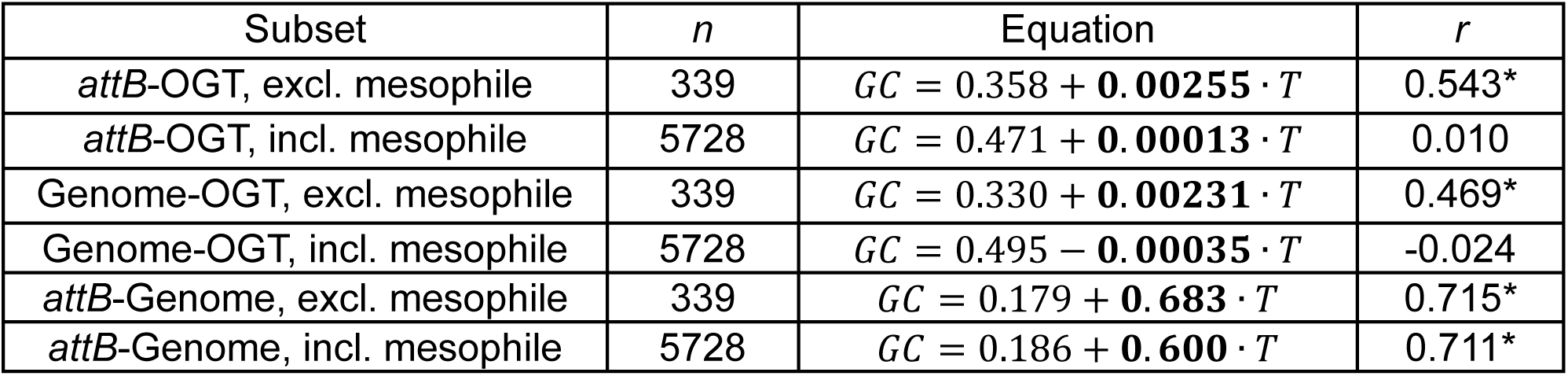
Analysis of tyrosine integrase-associated *attB* and genome-wide GC content. Integrases were separately analyzed based on their family (serine or tyrosine) and with the exclusion or inclusion of mesophile-class annotations. Equations represent calculated linear relationships between temperature and *attB* GC content. Tyrosine integrase-associated *attB* sites showed a statistically significant positive relationship to OGT when mesophile entries were excluded (*r*, Pearson correlation coefficient), whereas serine integrase-associated *attB* sites did not show statistically significant positive trends (**Supp. Table S1**). Tyrosine integrase-associated genome-wide GC content also showed a statistically significant positive relationship to OGT when mesophiles were excluded. Tyrosine integrase-associated *attB* sites and genome-wide GC content showed a moderately positive and statistically significant relationship in all analyses. Starred values indicate statistical significance (P < 0.05).

Whole-genome GC content and 16S rRNA gene GC content were compared with host OGT where available to assess whether *attB* GC trends reflected broader genome-wide nucleotide composition (**Supp. Fig. S1N**).

### Integrase clustering and thermal-class mixing analysis

To compare temperature homogeneity across integrase protein clusters, the clustering software MMSeqs2 (32) was used to cluster 227,967 unique integrase protein sequences (24), from hosts with known optimal growth temperatures, at 30%, 50%, and 70% amino acid identity (with a length coverage of 80%) to determine whether homologous integrases were found in hosts from multiple temperature classes. Five temperature classes were considered: psychrophile, psychrotroph, mesophile, thermophile, and hyperthermophile. Cluster purity was defined as the fraction of cluster members assigned to the dominant thermal class. Clusters with purity below 0.9 were defined as substantially mixed. Trees were also generated using RAxML-NG (33) to compare integrase similarity (For tree availability, see **Data Availability Statement; Supp. Fig. S2).**

### Design of reporter and integrase plasmids

Reporter plasmids (pRep; **Supp. Table S2**) were designed to contain two markers that were utilized for determining integrase-mediated inversion in subsequent experiments: a *sacB* sucrose lethality gene marker (34) in the forward orientation and a *specR* resistance marker in the reverse orientation (*specR_inv*). Reporter plasmids also contain *attB* and *attP* landing sites in an antiparallel orientation that uniquely pair with each test integrase. A promoter from the 16S rRNA region of *Thermus thermophilus* was used as a constitutive promoter upstream of the *attP* site to drive expression of the selectable marker in either orientation, and a terminator was positioned downstream of the *attB* site to prevent read-through transcription. This cassette (*attP*-*sacB*-*specR_inv*-*attB*) was flanked by *BsaI* sites for Golden Gate assembly (35) of reporter plasmids. The backbone contained a *repE/incC* mini-F origin for plasmid maintenance at single copy levels. Additionally, a *cmR* selectable marker was constitutively expressed to confer chloramphenicol resistance.

An integrase plasmid (pInt) was designed to contain a cognate integrase that specifically catalyzed a reaction between the paired *attB* and *attP* found on the reporter plasmids or an RFP CDS as a negative control (pNeg). This integrase (or RFP) was controlled by the arabinose-inducible P_araBAD_ promoter (36) and which was repressed by *araC* on the same plasmid under the control of a constitutive promoter. This vector contained a relatively low copy (∼10-20 copies/cell) p15A origin of replication and carried a selectable *kanR* or *tetR* marker.

### Cloning of thermophile-and psychrophile-derived integrases in *E. coli*

To assemble integrase plasmids, a gBlock (IDT) containing a codon-optimized ΦC31 integrase sequence or an RFP sequence (IDT codon optimization tool; **Supp. Table S3**) was combined with a vector backbone containing p15A and *kanR* or *tetR*, to generate pJLC201 or pJLC235 (pInt), or pJLC199 or pJLC255 (pNeg) respectively, in a one pot Golden Gate reaction (35) as per NEB guidelines using the NEBridge BsaI-HFv2 Golden Gate Assembly Kit (NEB Cat. #R3733).

Transformations were performed using a standard methodology for transforming chemically competent cells. Briefly, following thermal cycling (37°C for 60 min, 65°C for 10 min, 4°C hold), 5-10 µL of reaction mixture was transferred to 50 µL of chemically competent *E. coli* NEB DH5α cells (Cat. #C2987) with minimal agitation, followed by a 30-minute incubation on ice. Cells were heat shocked at 42°C for 30 seconds, then returned to ice for an additional 5 minutes. The resulting transformation was recovered in 1mL of SOC media at 37°C for 1 hour followed by plating on appropriate antibiotic selection plates (kanamycin [30 µg/mL] or tetracycline [10 µg/mL]).

Colonies were screened for successful assembly using colony PCR with Q5 DNA polymerase (NEB Cat. #M0491). For colony PCR, single isolated colonies appearing on selective plates were resuspended in 20 µL of ddH_2_O and added to a 25 µL Q5 reaction containing appropriate screening primers (P140 and P902; **Supp. Table S4**). PCR reactions were run on an agarose gel (1-2%) and reactions showing the appropriate size bands were sequenced at plasmidsaurus *via* Oxford Nanopore long-read sequencing. The resulting plasmids (pInt plasmids pJLC201 or pJLC235, and pNeg plasmids pJLC199 or pJLC255, kanamycin or tetracycline resistant respectively; **Supp. Table S2**) were then used as a vector backbone for subsequent cloning of integrase or control protein-bearing plasmids.

Similar methodology was used to assemble extremophile integrase sequences into pJLC235. Briefly, extremophile integrases were codon optimized (IDT codon optimization tool) and resulting gBlocks were cloned using the above Golden Gate methodology into nested BsaI sites in pJLC235. Cells were transformed as above into NEB DH5α cells and plated on selective media (tetracycline [10 µg/mL]). Successful clones were identified *via* colony PCR and validated *via* nanopore sequencing.

Reporter plasmids were constructed with a similar approach, developing a paired plasmid for the ΦC31 integrase that contained cognate *attB* and *attP* sites flanking *sacB* and *specR* as described above. Separate gBlocks containing these elements (**Supp. Table S3**) were cloned into a modified pCC1BAC vector (37) containing the *repE/incC* mini-F origin for single copy plasmid maintenance using Golden Gate following the above protocol with BsmBI sites to generate pJLC198 (**Supp. Table S2**). The protocol followed above methodology, using the NEBridge BsaI-HFv2 Golden Gate Assembly Kit (NEB Cat. #E1602) and the multiple insert methodology (35). Reaction mixtures were transformed into chemically competent NEB DH5α cells as described before plating on selective media (chloramphenicol [30 µg/mL]). Successful clones were screened and validated for use in inversion assays. Additional integrase-specific reporter plasmids (**Supp. Table S2**) were generated using Golden Gate assembly to insert gBlocks containing complementary *att* sites into nested BsmBI sites in pJLC198 as described above.

For inversion assays, each validated integrase plasmid and complementary reporter plasmid was purified by the PureYield Midiprep system (Promega Cat. #A2492) following manufacturer instructions and co-transformed into chemically competent *E. coli* NEB Turbo (Cat. #C2984). Transformations were performed as described above. Briefly, 1µL of each plasmid was added to a 50µL aliquot of chemically competent NEB Turbo cells with minimal agitation followed by a 30-minute incubation on ice. Cells were heat shocked at 42°C for 30 seconds, then returned to ice for an additional 5 minutes. The resulting transformation was recovered in 1mL of SOC media at 37°C for 1 hour followed by plating on appropriate antibiotic selection plates (chloramphenicol [30 µg/mL] for reporter plasmid and kanamycin [30 µg/mL] or tetracycline [1 µg/mL] for the integrase plasmid for reported studies). Overnight transformation plating was conducted at 25°C for thermophile-derived candidate pairs and 37°C for psychrotroph-derived candidate pairs to avoid unintended or premature temperature mediated integrase activity.

*E. coli* NEB Turbo carries an F’ plasmid encoding the F pilus required for infection by F-specific phages such as MS2 (38). Because pRep shares the same origin of replication as the native F’ plasmid, we used loss of MS2 susceptibility to confirm displacement of the F’ plasmid after pRep transformation (**Supp. Fig. S3**). Thus, we do not expect confounding effects from incompatible plasmid maintenance.

### Testing temperature-induced integrase expression using inversion assays in *E. coli*

To maintain consistent temperature profiles for inversion assays, outgrowth was conducted at 25°C, 30°C, 37°C or 42°C in LB Broth picking a single colony from overnight transformation plates of each integrase/reporter pairing with appropriate antibiotic selection (kanamycin/tetracycline and chloramphenicol at above concentrations). Integrases tested here are expressed under the control of an arabinose inducible promoter (P_araBAD_) and were therefore induced with 0.2% arabinose or quenched with 0.2% glucose and grown in 5mL LB Broth (Lennox; Sigma Cat. #L3022) containing appropriate antibiotic selection and induction media overnight at each temperature. At stationary phase, overnight cultures were serially diluted 10-fold in SM buffer (Teknova Cat. #S0249) ranging from 0 to 10^-7^. For all dilutions, 2.5 µL of cells were spotted in technical triplicate on LB with 1.5% w/v agar (Fisher, Cat. # BP97445) containing appropriate antibiotics in the absence or presence of 5% sucrose (Sigma Cat. #S5391). The addition of 5% sucrose proves lethal to *E. coli* cells expressing *sacB* and can be used to assess integrase mediated flipping of the *attP*-*sacB*-*specR_inv*-*attB* cassette (34). Likewise, spectinomycin [50 µg/mL] was used to assay for integrase activity during assay development. Plated cells were incubated overnight at 25°C, 30°C, 37°C or 42°C, corresponding to the temperature of overnight culture. After 24-36 hours, the resulting colonies were counted and select colonies were picked for colony PCR using Q5 DNA polymerase and sequenced to confirm inversion events.

For amplicon sequencing, PCR products were verified for size and quantified, then submitted to Plasmidsaurus (Eugene, OR) for sequencing. Samples were sequenced on Oxford Nanopore Technologies (ONT) R10.4.1 flow cells using a proprietary primer-free protocol.

### Quantification of integrase-mediated inversion activity

Integrase-mediated inversion activity was quantified by normalizing colony recovery on sucrose counterselection plates to the average colony recovery on growth-control plates per biological replicate, selecting only for maintenance of the two-plasmid system. For each condition, normalized integrase activity was calculated as:

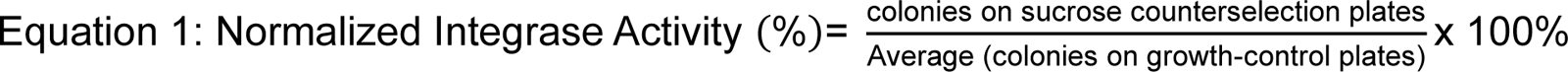

Each assay condition was performed with three independent biological replicates, and each biological replicate was plated in three technical replicate spots. Technical replicate counts were averaged within each biological replicate prior to statistical analysis, such that statistical comparisons were performed using n=3 independent biological replicates per condition. For Log-scale graphs, a limit of detection (LOD) was imposed at a Normalized Integrase Activity value of 0.0001%. Activity values were calculated from colony counts obtained from serial dilution spot plates and were used to compare recombination activity across induction conditions, temperatures, and integrase candidates.

### Statistical analysis

Statistical analyses were performed in GraphPad Prism. Unless otherwise stated, each condition included at least three independent biological replicates, each plated in at least two technical replicate spots. Technical replicate colony counts were averaged within each biological replicate before statistical testing; therefore, statistical analyses were performed using n≥3 independent biological replicates per condition.

Comparisons between two groups were performed using a lognormal Welch’s t-test. Comparisons among multiple temperature or induction conditions were performed using ordinary one-way ANOVA as indicated in the figure captions

## RESULTS

### Linking integrases to host thermal niche

Database integration as described above (see **Materials and Methods**) generated a curated database of 12,760 genome assemblies with associated temperature annotations (**Figure 1A**), providing the foundation for mapping integrase and attachment (*att*) site sequences to host growth-temperature profiles. Although more than twelve thousand entries are represented in the database, the dataset is strongly biased toward mesophiles which was considered throughout the following analysis. When we quantified the number of entries represented at each reported OGT, we observed clear enrichment at 37°C (**Supp. Fig. S1M**). This enrichment may reflect bias toward convenience of culturing at 37°C more than the true OGT for all organisms.

**Figure 1.**
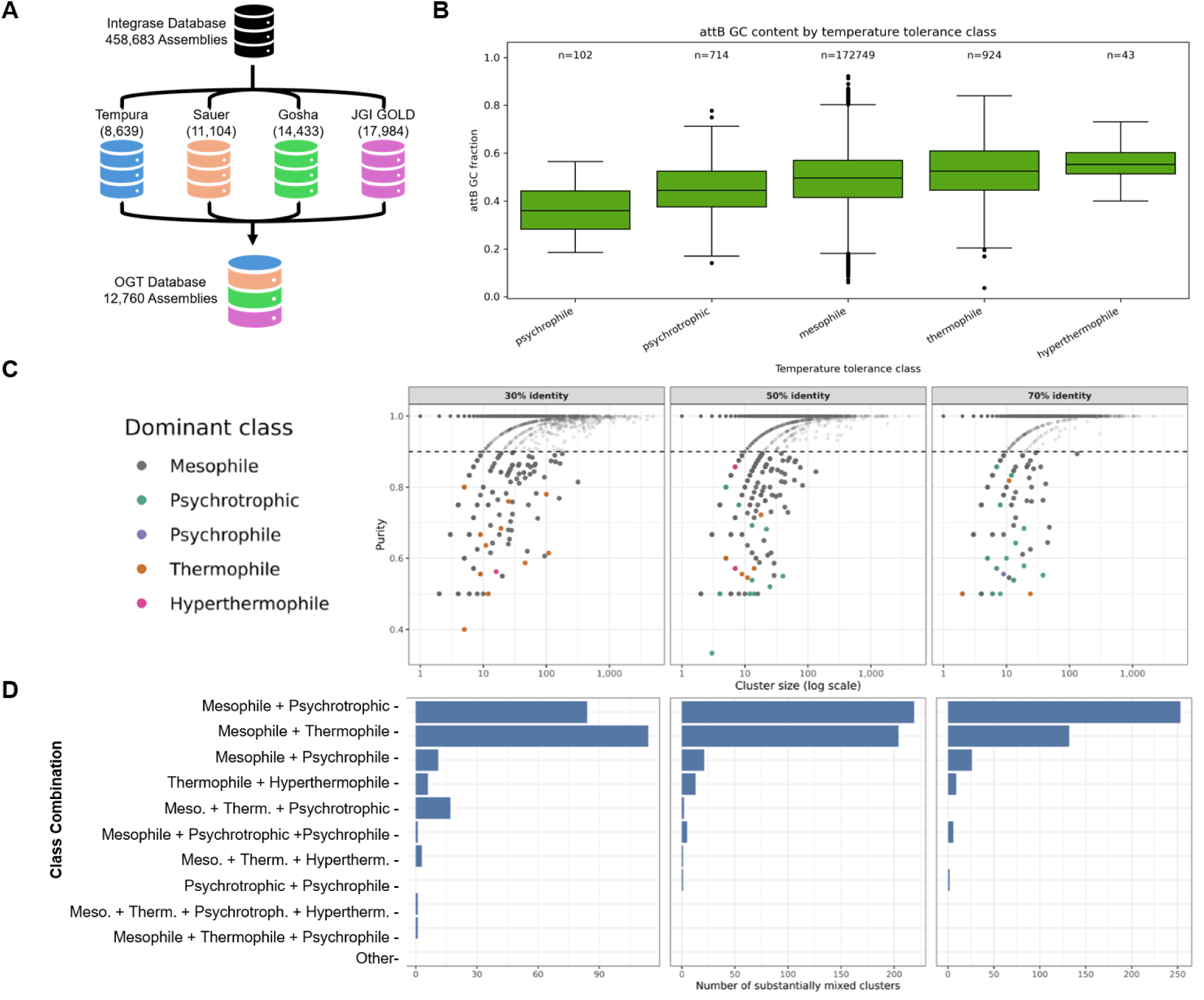
Large-scale genomic integrase mining reveals temperature-dependent trends. **(A)** Growth temperature information from four datasets (see **Materials** for descriptions) was available for 12,760 of the 458,683 genomes underlying the TIGER/Islander integrase/*att* database (24). **(B)** GC content of integrase-associated *attB* sites separated by temperature class **(C)** Number of clusters at 30%, 50% and 70% identity thresholds plotted by temperature purity by number of cluster members. Clusters with less than 90% purity fall below the dashed line and are colored by their dominant temperature type. **(D)** Combination of temperature types present in substantially mixed clusters (those at or below 90% purity) by each clustering identity thresholds.

### GC Content of tyrosine-associated *attB* sites increases with host growth temperature

To investigate sequence-level features associated with host thermal environments, we analyzed the GC content of *attB* sites corresponding to integrases across different temperature classes. When we linked temperature annotations to *attB* sites from Integrases-on-Demand (29), a modest trend was observed in which *attB* GC content increased progressively from psychrophilic to hyperthermophilic hosts (**Figure 1B**). Serine and tyrosine integrases were analyzed separately because they represent evolutionarily and mechanistically distinct recombinase families (**Supp. Table S1**). We also evaluated this relationship excluding mesophile-annotated hosts to avoid potential sampling and culturing bias effects. We show statistically significant relationship between host growth temperature and *attB* GC content for tyrosine integrase *attB* sites, when mesophiles were excluded from analysis (**Table 1; Supp. Fig. S1A-B**). Serine integrase *attB* sites did not show a significant temperature-GC correlation which may result from a small sample size, especially for lower reported OGT (**Supp. Fig. S1G-L)**. Additionally, a lack of significant relationships when mesophile populations were included indicates that bias from experimental culturing conditions may mask temperature-associated trends in large datasets.

This pattern is consistent with known relationships between GC content and nucleic acid stability at elevated temperatures, which suggest that high-GC *att* sites may reduce recombination efficiency at low temperatures by increasing the energetic cost of DNA melting or junction formation (39). Likewise, high-temperature hosts impose additional selection for thermostable integrases (12). Thus, thermal niche may shape both *att*-site composition and integrase stability. Our observation of an *attB* GC-OGT relationship may not be a simple by-product of genome-wide GC content trends. Prior work reports a positive relationship between genome-wide GC content and OGT (40), and our analysis also detected a positive correlation between genome-wide GC and OGT when mesophile genomes are excluded, as well as a broad correlation between *attB* and genomic GC across integrase subsets (**Supp. Fig. S1C-F**). A Steiger’s Z-test to compare the correlation of tyrosine *attB* and genome-wide GC content excluding mesophiles showed a statistically significant increase in the Pearson correlation coefficient for the *attB* GC vs genome-wide GC (Z = 2.13, P = 0.016). Further, we observed that GC content of the 16S rRNA gene is linked to increasing OGT in the TEMPURA database, whereas genome-wide GC content did not appear to have an obvious trend when analyzed on a per-database basis (**Supp. Fig. S1N**).

### Integrases sequences cluster based on host thermal class

We found that homology-based clustering of integrase proteins showed that similar integrases tend to originate from hosts occupying similar thermal niches. The 227,967 unique integrase protein sequences from temperature-annotated hosts in TIGER/Islander (24) were predominantly from mesophiles with 225,149 integrase sequences (98.76%), followed by thermophiles with 1,451 (0.64%), psychrotrophs with 1,187 (0.52%), psychrophiles with 113 (0.05%), and hyperthermophiles for with (0.03%).

Despite this imbalance, temperature mixing across integrase clusters was relatively uncommon and decreased with increasing clustering stringency. Mixed clusters represented 12.5% of clusters at 30% amino acid identity, 4.0% at 50% identity, and 1.2% at 70% identity. When only substantially mixed clusters were considered, defined as clusters with less than 90% thermal-class purity, the fraction was lower: 4.2% at 30% identity, 2.2% at 50% identity, and 0.94% at 70% identity (**Figure 1C**).

When we examined which thermal classes co-occurred within substantially mixed clusters, we found that mesophile–extremophile mixtures were the most common mixed-cluster type. Clusters containing both hot-associated and cold-associated classes were not observed at all at 50% or 70% amino acid identity (**Figure 1D**) and were rare even using a loose 30% threshold. Thus, while broad integrase families can include members from multiple thermal classes, finer-grained clusters were usually temperature-homogeneous.

### Selection of candidate integrases from temperature-annotated hosts

Based on the temperature-linked integrase database, we selected representative integrase and *att*-site pairs for experimental validation. Candidates were prioritized from organisms with high-confidence temperature annotations and available predicted cognate *attB* and *attP* sites, enabling direct construction of matched reporter and integrase plasmids. We focused on candidates spanning distinct thermal niches, including thermophile-derived integrases from *Thermus thermophilus* (GCA_000091545.1) and *Geobacillus stearothermophilus* (GCA_003667675.1), as well as a lower-temperature-associated integrase from the psychrotroph *Pseudomonas cerasi* (GCA_014450905.1) (**Table 2**). The ΦC31 integrase was included as a non-temperature-selected benchmark expected to exhibit robust recombination across the tested *E. coli*-compatible temperature range (1,41). This candidate set allowed us to test whether host thermal niche corresponded to temperature-dependent recombination activity in a controlled inversion assay.

**Table 2:** Candidate integrases for experimental validation of temperature-dependent recombination. Integrases were selected from temperature-annotated hosts to probe thermal switching from cold-to hot-associated OGTs. The ΦC31 integrase is expected to show robust activity across the range of compatible *E. coli* temperatures. Thermophile integrase candidates from *T. thermophilus* and *G. stearothermophilus* were selected as they comprise well-studied extremophiles. The *P. cerasi* integrase represents a psychrotroph-derived candidate with predicted functionality at lower temperatures. Genome assembly IDs were obtained from the TIGER/Islander (24) database. OGT annotations were obtained as described **(Methods; Figure 1A**).

| Candidate Source | Identifier | Expected phenotype | OGT |
| --- | --- | --- | --- |
| ΦC31 | GCA_000848045.2 | Non-switching benchmark | 37°C |
| *T. thermophilus* | GCA_000091545.1 | hot-ON/cold-OFF | 67°C |
| *G. stearothermophilus* Gs1 | GCA_003667675.1 | hot-ON/cold-OFF | 55°C |
| *G. stearothermophilus* Gs2 | GCA_003667675.1 | hot-ON/cold-OFF | 55°C |
| *P. cerasi* | GCA_014450905.1 | cold-ON/hot-OFF | 28°C |

### Toxin inversion assay enables quantitative measurement of integrase-mediated recombination

To establish a quantitative assay for integrase activity in *E. coli*, we first designed integrase-expression and reporter plasmids for the benchmark ΦC31 integrase, which is expected to show robust recombination across the tested temperature range. The integrase plasmid, pInt, encoded ΦC31 integrase under control of the arabinose-inducible P_araBAD_ promoter as described above (see **Materials and Methods**). The reporter plasmid, pRep, contained a recombination cassette in which ΦC31 *attP* and *attB* sites were positioned in opposing orientations around a *sacB* counterselection marker and an inverted specR antibiotic resistance marker. This cassette, *attP*-*sacB-specR_inverted_*-*attB*, was positioned downstream of a constitutive promoter (P_16S_; **Supp. Table S3**), such that *sacB* is expressed before inversion and *specR* is expressed after inversion (**Figure 2A**).

**Figure 2.**
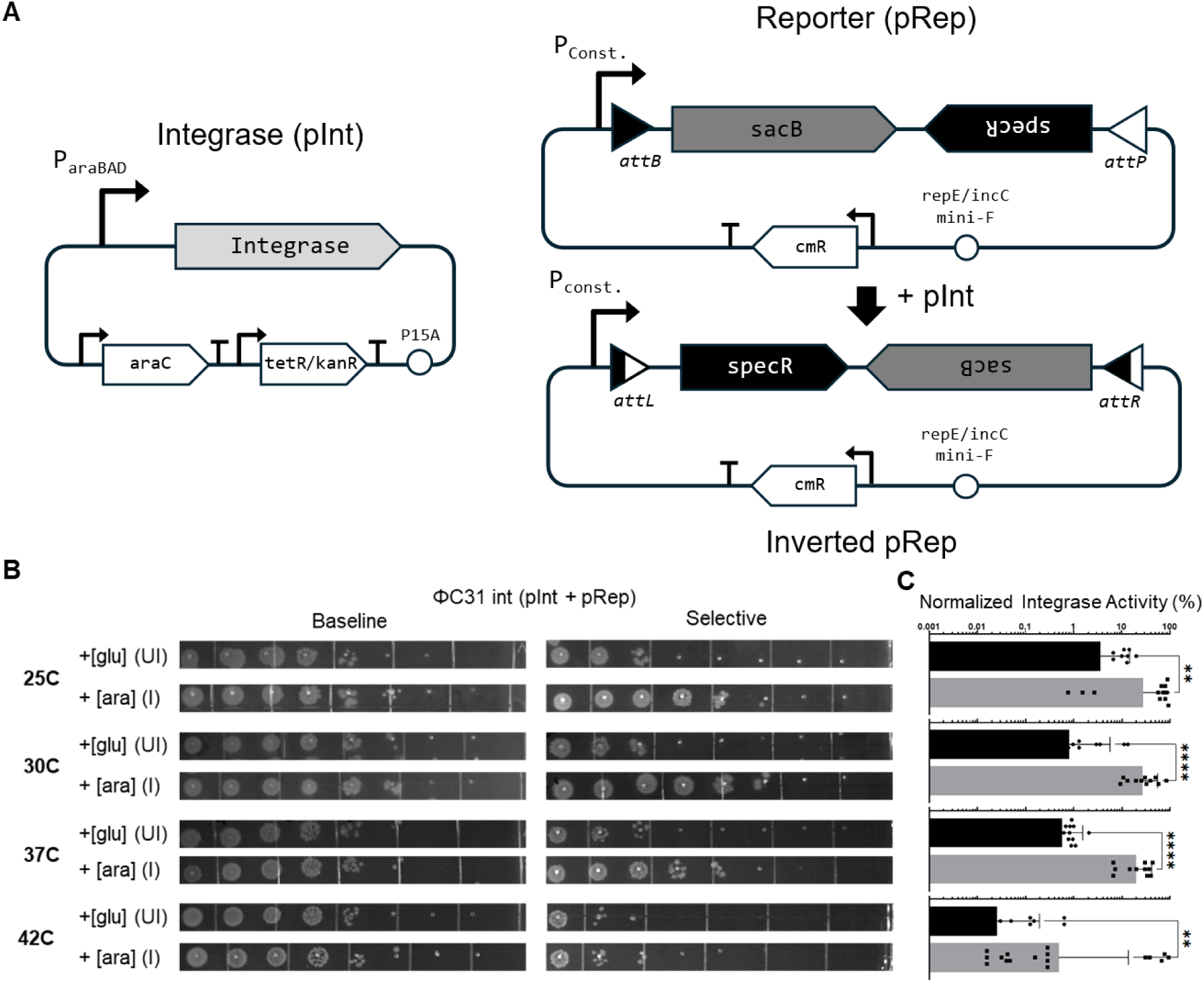
Integrase-mediated inversion testbed development. A two-plasmid system **(A)** was used for assaying extremophile integrase activity. In the reporter plasmid (pRep; **Supp. Table S2**), a strong constitutive promoter (P_16s_) controls *sacB*-mediated counterselection pre-inversion. This cassette is flanked by *attB* and *attP* sites which pair with a cognate integrase on a second plasmid (pInt; **Supp. Table S2**) under the control of an inducible promoter (P_araBAD_). Serial dilutions were plated **(B)** and quantified **(C)** to assess integrase activity for the ΦC31 integrase and *att* pair. Integrase production was induced (arabinose, 0.2% w/v) or quenched (glucose, 0.2% w/v) and cells were plated on plates containing 5% w/v sucrose to select for inversion events or on plates that just contained selection for the two-plasmid system (Cm/Tet). RFP was used as a negative control in place of the ΦC31 integrase (pNeg; **Supp. Fig. S4**; **Supp. Table S2**) to check for endogenous integrase-mediated inversion events. For statistical analysis a Welch’s t-test was performed in GraphPad Prism. *, P<0.05; **, P<0.01; ***, P<0.005; ****, P<0.0005.

Upon induction of integrase expression with arabinose, ΦC31 catalyzes recombination between *attP* and *attB*, inverting the cassette and placing *sacB* in the antisense orientation relative to the upstream promoter. Although *specR* was included as an orthogonal post-inversion marker, we did not use spectinomycin selection as the primary quantitative readout because each candidate integrase carries distinct *attP* and *attB* sequences, and recombination generates candidate-specific *attL* and *attR* junctions. These variable *att*-site sequences could differentially affect *specR* expression by introducing cryptic promoter or terminator-like elements, confounding the assay. We therefore used *sacB*-based sucrose counterselection as the primary phenotypic readout for inversion (**Figure 2B**).

Integrase activity was quantified as the ratio of colonies recovered on sucrose counterselection plates to colonies recovered on growth-control plates selecting only for maintenance of the two-plasmid system (**Eq. 1**). This normalization allowed recombination-associated colony recovery to be compared across temperatures and induction conditions while accounting for differences in recovery of viable two-plasmid-containing cells.

The benchmark ΦC31 integrase showed robust inversion activity across the tested temperature range, with no significant temperature-dependent change in normalized activity (**Figure 2C**). In contrast, induction state strongly affected recombination: arabinose-induced cultures showed significantly higher integrase activity than glucose-repressed cultures. When RFP was expressed in place of ΦC31 integrase (pNeg), integrase activity was below the limit of detection (**Supp. Fig. S4A–C**).

To ensure that sucrose-resistant recovery reflected programmed cassette inversion rather than nonspecific escape from *sacB* counterselection, we validated representative colonies by colony PCR and amplicon sequencing. These analyses confirmed the expected post-recombination reporter architecture, including diagnostic inversion junctions, in colonies recovered from both the ΦC31 benchmark assay and candidate integrase assays. This molecular confirmation supports normalized sucrose-resistant recovery as a quantitative phenotypic readout of integrase-mediated inversion.

### Thermophile-derived integrases exhibit hot-ON/cold-OFF recombination activity

To test whether a thermophile-derived integrase exhibits temperature-dependent recombination in the inversion assay, we first constructed matched reporter and integrase-expression plasmids for a candidate integrase from *T. thermophilus* (**Table 2; Supp. Table S2**). As with the ΦC31 integrase, pInt and pRep plasmids or pNeg and pRep were co-transformed and integrase-mediated inversion was assayed at test temperatures ranging from 25°C to 42°C. For this thermophile-derived integrase candidate, co-transformations were plated at 25°C to prevent premature flipping events. Following induction or quenching, overnight cultures of cells at stationary phase were serially diluted and plated on selective media (5% sucrose) to assess for effective inversion of the *sacB*-encoded toxin (**Figure 3A**).

**Figure 3.**
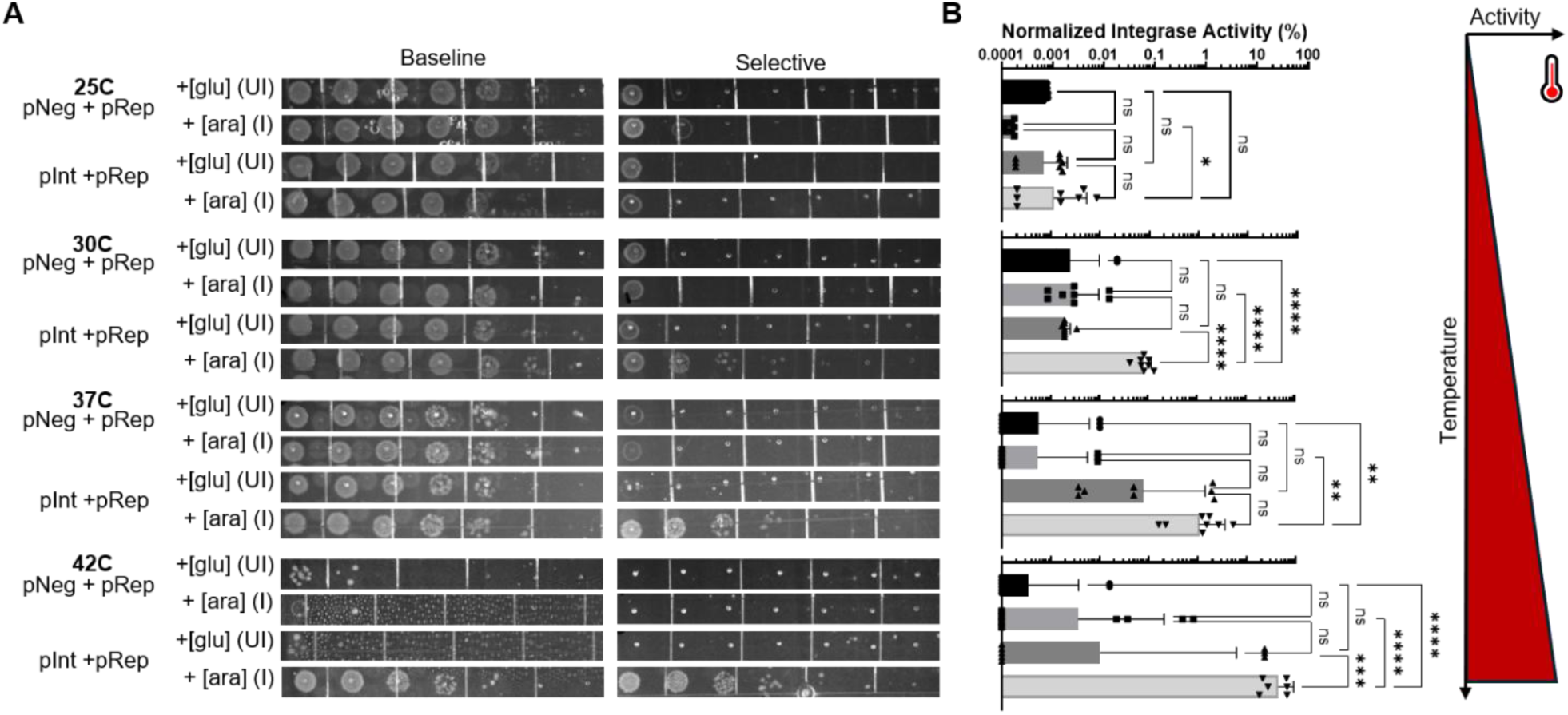
Inversion assay of thermophile *Thermus thermophilus-*derived integrase in *E. coli*. Serial dilutions were plated **(A)** and quantified **(B)** to assess integrase activity for the *T. thermophilus*-derived integrase and *att* pair at four different temperatures (25°C, 30°C, 37°C, and 42°C). Integrase production was induced (arabinose, 0.2% w/v) or quenched (glucose, 0.2% w/v) and cells were plated on plates containing 5% w/v sucrose to select for inversion events or on plates that just contained selection for the two-plasmid system (Cm/Tet). RFP was used as a negative control in place of the *T. thermophilus* integrase to check for endogenous integrase-mediated inversion events. For statistical analysis an ordinary one-way ANOVA was performed in GraphPad Prism. *, P<0.05; **, P<0.01; ***, P<0.001; ****, P <0.0001.

We show that a candidate *T. thermophilus* derived integrase (pInt + pRep) has significantly increased activity above the negative control (pNeg + pRep) and likewise shows increased activity when induced (0.2% w/v arabinose) as compared to quenched (0.2% w/v glucose) conditions, although leaky integrase expression was observed in all conditions. Moreover, we show that this activity is dependent on temperature, with the highest integrase activity at 37°C and 42°C (**Figure 3B**). These findings are consistent with our original hypothesis that temperature can be harnessed as an intrinsic biological control over extremophile-derived integrase activity.

To test a broader repertoire of thermophile-derived integrases, we further tested integrases from two strains of *Geobacillius stearothermophilus* (**Supp. Fig. S5A-B**). While both showed increased integrase activity with increasing temperature, *G. stearothermophilus* Gs1 followed a similar pattern of increasing activity when approaching the maximum stable growth temperatures of *E. coli* (∼42°C) (**Supp. Fig. S5A**), *G. stearothermophilus* Gs2 showed decreased activity only at 25°C (**Supp. Fig. S5B**), indicating a potentially lower thermal switching temperature that merits further investigation.

Interestingly, for thermophile candidates, we observed varying degrees of endogenous *E. coli* integrase activity across all temperatures. While candidate pairings (pInt + pRep) were significantly higher than integrase-free controls (pNeg + pRep), inversion events were nonetheless observed in groups lacking the *T. thermophilus*-derived integrase (**Figure 3A-B**), and even for *E. coli* containing pRep alone (**Supp. Fig. S6A-C**). Furthermore, *G. stearothermophilus att* pairings showed increased endogenous *E. coli* integrase activity (**Supp. Fig. S5A-B**), as compared to *T. thermophilus*-derived *att* sites. These results suggest that a subset of heterologous *att*-site pairs may be susceptible to integrase-independent background inversion in the *E. coli* host context.

### A psychrotroph-derived integrase shows lower-temperature-restricted activity

To determine whether integrases from lower-temperature-associated hosts can exhibit temperature-response profiles distinct from thermophile-derived candidates, we tested an integrase and predicted cognate *att*-site pair from the psychrotroph *P. cerasi* (**Table 2; Supp. Table S3**). Although *P. cerasi* is not an extreme psychrophile, its reported growth range lies near the lower end of the mesophilic range, making it compatible with an *E. coli* testbed while still allowing us to ask whether lower-temperature-associated hosts can yield integrases with activity biased toward cooler conditions. Experiments were performed with *P. cerasi* integrase, cognate *att* sites and controls as described above tested across 25°C, 30°C, 37°C, and 42°C using sucrose counterselection as the primary readout for reporter inversion.

The *P. cerasi*-derived integrase displayed a temperature-response profile distinct from the thermophile-derived integrases. In arabinose-induced cultures, recombination-associated colony recovery was highest at lower temperatures and not detected above background at elevated temperatures, indicating a cold-ON/hot-OFF activity profile (**Figure 4A-B**). Interestingly, the highest reported activity was shown at 30°C when induced. Although the absolute activity of the *P. cerasi* integrase was lower than that of the benchmark ΦC31 system, the lower-temperature signal was reproducible and distinguishable from no-integrase controls under the tested conditions.

**Figure 4.**
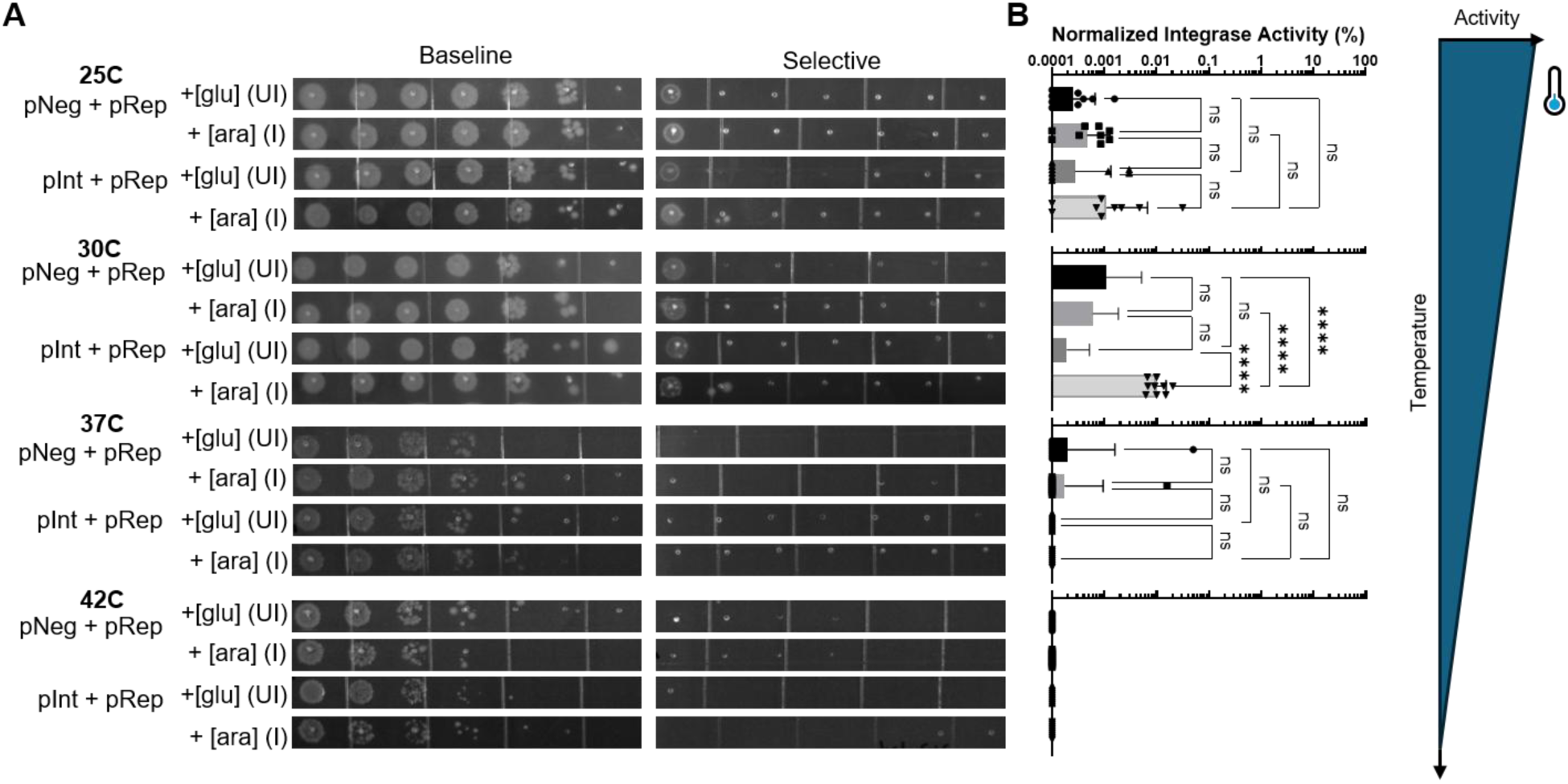
Inversion assay of *Pseudomonas cerasi-*derived integrase in *E. coli*. Serial dilutions were plated **(A)** and quantified **(B)** to assess integrase activity for the *P. cerasi* integrase and *att* pair at four different temperatures (25°C, 30°C, 37°C, and 42°C). Integrase production was induced (arabinose, 0.2% w/v) or quenched (glucose, 0.2% w/v) and cells were plated on plates containing 5% w/v sucrose to select for inversion events or on plates that just contained selection for the two-plasmid system (Cm/Tet). RFP was used as a negative control in place of the *P. cerasi* integrase to check for endogenous integrase-mediated inversion events. For statistical analysis an ordinary one-way ANOVA was performed in GraphPad Prism. *, P<0.05; **, P<0.01; ***, P<0.001; ****, P <0.0001.

This lower-temperature-restricted profile suggests that temperature-responsive integrases may be discoverable not only from classical thermophiles, but also from organisms occupying cooler or lower-mesophilic niches. The *P. cerasi* candidate therefore complements the thermophile-derived candidates by demonstrating that temperature-biased integrase activity can occur in both directions across the tested thermal range.

### Host thermal niche is associated with integrase temperature-response profiles

To further determine whether temperature-response profiles were associated with host thermal niche we performed a statistical comparison of induced recombination activity across all tested integrases. The benchmark ΦC31 integrase was active across the tested temperature range and did not show a statistically significant temperature-dependent shift in normalized recombination activity (**Figure 5**). However, the 42°C ΦC31 condition showed a broader, apparently bimodal distribution that was not observed for the other integrase systems tested at 42°C. Because ΦC31 is a non-thermophile-associated *Streptomyces* phage system and 42°C approaches the upper practical range of this *E. coli* assay, this pattern may reflect component-specific instability or threshold behavior of the ΦC31 integrase-*att* reporter system under brittle assay conditions rather than a bona fide temperature-switching phenotype.

**Figure 5.**
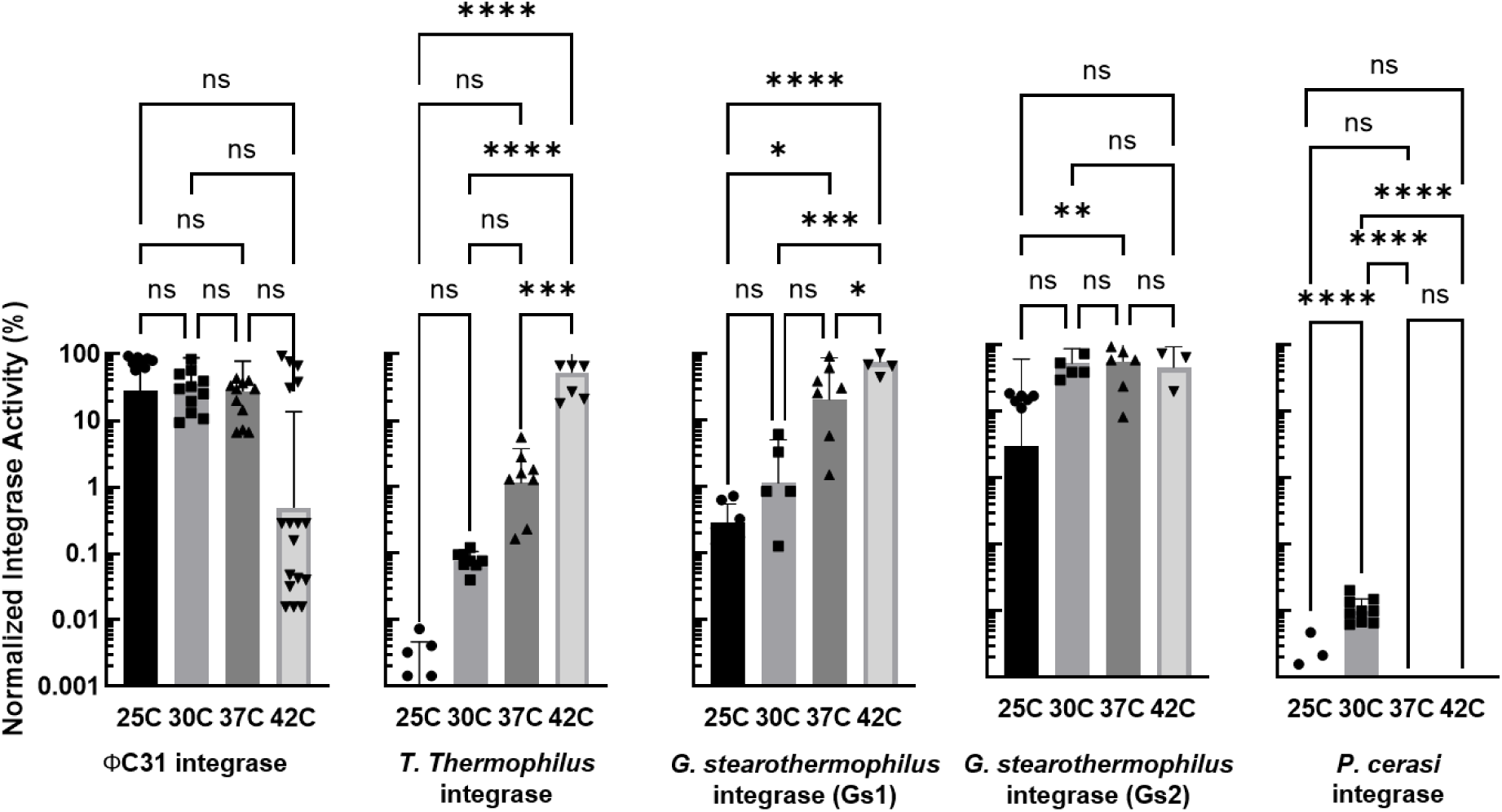
Extremophile-derived integrases can exhibit temperature switchability. Quantification of integrase activity when induced (arabinose, 0.2% w/v). Activity is compared at four temperatures (25°C, 30°C, 37°C, and 42°C) for five integrases derived from ΦC31, *T. thermophilus*, *G. stearothermophilus* Gs1*, G. stearothermophilus* Gs2, and *P. cerasi*. Cells were plated on plates containing 5% w/v sucrose to select for inversion events or on plates that just contained selection for the two-plasmid system (Cm/Tet). For statistical analysis an ordinary one-way ANOVA was performed in GraphPad Prism. *, P<0.05; **, P<0.01; ***, P<0.001; ****, P <0.0001.

In contrast, thermophile-derived candidates showed hot-biased activity, although the strength and apparent transition temperature differed among integrases. The *T. thermophilus* and *G. stearothermophilus* Gs1 integrases showed significantly higher activity at 42°C than at lower assay temperatures, consistent with hot-biased recombination. The *G. stearothermophilus* Gs2 candidate showed a weaker temperature response, with a significant difference detected between 25°C and 37°C, suggesting a lower or broader activity transition.

The *P. cerasi* integrase displayed the opposite pattern. Its activity was maximal at 30°C, and all comparisons to the 30°C condition were significant. Activity at 25°C was detectable but near baseline, and differences between 25°C and the other elevated-temperature conditions were not significant. No detectable activity was observed at 37°C or 42°C. Together, these statistical comparisons support the central prediction that host thermal niche can guide discovery of integrases with distinct temperature-response profiles.

## DISCUSSION

### Temperature-linked genomes are a discovery engine for intrinsic thermal control

While the literature is rich in examples of temperature-sensitive enzymes, a lack of systematic roadmaps limits discovery throughput (11,12,42). Here, we linked prokaryotic genomes to curated growth-temperature metadata and connected those thermal annotations to integrase sequence and their cognate attachment sites. Likewise, integrase function depends on both the integrase protein and the interaction with its cognate *attP* and *attB* sites (1,17,43). We utilized the TIGER/Islander database to map integrases to their predicted attachment sites, turning genome mining into experimentally testable recombinase systems (24).

The broader value of our work is not limited to integrases. This study establishes a way to search biology by thermal niche. A temperature-linked genome resource can be used to search for any biological component where intrinsic thermal control would be useful. Future studies could efficiently mine Cas systems, nucleases, polymerases, transcriptional regulators, metabolic enzymes, toxins, antitoxins, secretion systems, and other bioproducts where temperature-dependent activity matters.

### Integrase-*att* pairs carry thermal-niche structure

These computational analyses show that closely related integrases tended to come from hosts with similar thermal profiles, especially at higher clustering stringency. Mixed-temperature integrase clusters became rare as sequence identity increased. This suggests that integrase sequence similarity can help prioritize candidates from specific host temperature ranges.

Tyrosine integrase-associated *attB* sites showed increased GC content with host growth temperature, whereas serine integrase-associated *attB* sites did not show the same relationship. However, the tyrosine *attB* trend was apparent only after mesophile-class entries were removed, suggesting that the signal may be obscured by the large and potentially biased mesophile dataset, particularly the frequent annotation of 37°C as an optimal growth temperature. The absence of a comparable trend among serine integrase *attB* sites should also be interpreted cautiously, as serine integrase systems were much less represented in the dataset. Despite these limitations, the *attB* result is important because it suggests that temperature-responsive recombination may not be determined by integrase protein sequence alone. Attachment-site composition, including GC content and associated DNA melting behavior, may also influence recombination efficiency across temperatures. Thus, engineering temperature-switchable integrases may require tuning both the recombinase and its cognate *att* sites.

### Extremophile-derived integrases exhibit thermal switching

Our results support that integrases derived from extremophile hosts exhibit temperature-biased activity. Thermophile-derived integrases from *Thermus thermophilus* and *Geobacillus stearothermophilus* showed increased recombination frequency at higher temperatures, but differed in activity, transition behavior, and dynamic range. These differences are useful because they define an engineering search space rather than a single fixed phenotype.

Conversely, the *Pseudomonas cerasi* integrase showed the opposite trend. While not broadly cold-active, it is better described as lower-temperature-restricted, with an optimum activity near 30°C. Activity at 25°C was detectable but generally low, and differences between 25°C and the 37°C and 42°C temperature conditions were not statistically significant. Its absolute activity in *E. coli* was low compared with ΦC31, but the phenotype is still valuable, and activity may improve in other host contexts. Activity near 30°C and lack of detectable activity at 37°C or above make it a strong starting point for temperature-gated DNA-state control at reduced temperatures.

Together, these findings establish naturally occurring integrase diversity as a source of temperature-responsive recombinases and provide a foundation for engineering sharper, higher-activity temperature-switchable genetic control elements.

### Native extremophile integrases are a basis for temperature-dependent editors

The extremophile-derived integrases identified here establish a framework for temperature-biased recombination activity, but their switching properties will require refinement before deployment. Useful temperature-responsive integrase devices will need sharper transition thresholds, reduced off-state activity, increased on-state activity, and transition temperatures matched to specific applications. For example, a switch designed for 42°C bioprocess control would require different performance characteristics than one intended for fever-range activation, and a biocontainment switch designed to limit environmental release would impose still different requirements.

The *P. cerasi*-derived integrase presents a distinct engineering challenge because its on-state activity in *E. coli* was comparatively low. Improving this candidate would likely require increasing activity at the desired permissive temperature while preserving reduced activity at elevated temperatures. More broadly, the assay developed here provides a practical platform for such optimization. Normalized colony recovery provides a quantitative phenotype linked to integrase-mediated inversion, making both selection and counterselection strategies feasible. Variant libraries could be enriched at desired on-temperatures and depleted at undesired off-temperatures through iterative screening. Repeated rounds of selection could then tune the switching threshold, dynamic range, and leakiness of candidate integrases, providing a path from natural thermal bias to engineered thermal control.

### Temperature-based activity depends on host context

All candidates were tested in *E. coli* for proof-of concept due to its tractability and low cost for testing. Activity profiles in vivo will be subject to folding, expression burden, codon context, missing host factors, or host physiology. An extremophile-derived integrase may behave differently in its native background. The same enzyme may also show different profiles in probiotic strains, industrial microbes, environmental isolates, yeast, or human cells. Before these systems become general tools, they should be tested across host contexts, both before and after engineering.

Cell-free systems should be a useful next step. A TXTL-style assay could measure integrase activity across a continuous temperature gradient without confounding effects from growth, plasmid maintenance, or viability (44,45). Reactions could be supplemented with lysates from the native or thermally matched host to test whether missing host components limit activity in *E. coli*. Purified integrases produced under different temperature regimes could then help determine whether thermal gating reflects folding, stability, or conformational effects (46). These approaches would separate protein-intrinsic temperature dependence from host-dependent effects and generate higher-resolution switch curves.

### Background recombination defines an off-state design constraint

A key design constraint revealed by this study was background inversion in some reporter constructs. For several thermophile-derived reporter plasmids, sucrose-resistant colonies were recovered even in the absence of the cognate integrase-expression plasmid, and molecular analysis confirmed inversion of the reporter cassette. This background was generally low, with the highest reported activity at 5.42% (*G. stearothermophilus* Gs2; **Supp. Fig S5B**), and was not observed uniformly across all reporters, indicating that it is not an inherent property of the assay. Instead, it appears to depend on the specific *att*-site pair, local reporter sequence context, or interaction between the reporter and the *E. coli* host.

This observation is consistent with a broader feature of integrase biology. Integrase-*att* recognition is often stringent at the crossover core, but the flanking binding arms can tolerate sequence variation in some systems (1,17). This tolerance can broaden functional site recognition, allowing related or partially compatible sequences to participate in recombination at low frequency. Such behavior has been examined in limited contexts, but not systematically across the broad diversity of integrases and attachment sites now available through genome-scale resources. Our results suggest that host-dependent activity on heterologous *att* sites should be treated as a measurable property of each integrase-*att*, not merely assay noise.

The specific molecular source of the background remains unresolved. Native recombination pathways, cryptic site-specific recombinases, or resident prophage-derived functions could contribute. However, prophage genes are often transcriptionally silent under standard lysogenic conditions, as illustrated by lambda-like systems in which most phage functions are repressed unless the prophage is induced (47). Cryptic or degraded prophage remnants may also have altered regulatory logic or incomplete gene sets, making their behavior difficult to predict from annotation alone. Thus, if prophage-derived functions contribute to background inversion, their activity may depend strongly on host genotype, reporter sequence, and host physiology, including stress state or growth temperature. We evaluated background inversion in a prophage signature free strain of *E. coli* (48), and for the *T. thermophilus*-derived integrase observed no background integrase activity in cells with pRep and pNeg, but saw integrase activity when the *T. thermophilus* pInt and pRep were present (**data not shown**). Future experiments to probe the specific integrase(s) interacting with pRep in these conditions could be valuable for applications that rely on tight molecular control of integrase-*att* activity.

### Temperature can program genetic containment

Temperature-responsive integrases address a limitation that expression switches alone do not solve: control over genetic state. Many engineered microbes operate in environments with predictable temperature transitions, including host-associated microbes moving between ambient conditions and warm-bodied hosts, industrial cultures exposed to programmed temperature shifts, and environmental consortia experiencing daily or seasonal thermal cycles. These regimes can be treated not as passive constraints, but as programmable inputs for controlling DNA-state transitions.

One compelling application is temperature-programmed genetic self-removal (49,50). An engineered organism could execute a function under one thermal regime and excise the engineered cassette after transition to another, shifting containment from control of cell survival to control of genetic persistence. This distinction is important because cells, DNA, or mobile genetic elements may persist after the intended function is complete. A recombinase-based self-removal program directly targets the engineered modification. For example, a probiotic strain could activate a payload at host-associated temperature and remove the introduced cassette after exit into a cooler environment. Similar architectures could be adapted for microbiome engineering, environmental biosensing, bioproduction, or biological countermeasure delivery. Integrases are well suited for these designs because recombination creates durable DNA-state changes (18–20); depending on *att*-site arrangement and directionality control, the system can invert, integrate, excise, erase, or record genetic information.

### From thermal niche to biological control

This study makes host thermal niche a design variable for genome engineering. By pairing host growth-temperature data with mapped integrase-*att* systems, we identified natural recombinases with distinct temperature-response profiles, including hot-biased thermophile-derived integrases and a lower-temperature-restricted *P. cerasi* integrase. These findings show that thermal control can be discovered directly from natural diversity and converted into programmable DNA-state control. The approach should extend beyond integrases to Cas systems and other bioproducts where intrinsic temperature responsiveness would improve control, safety, and deployment.

## AUTHOR CONTRIBUTIONS

JHA: Conceptualization, Methodology, Validation, Analysis, Investigation, Data Curation, Visualization, Writing – original draft, Writing – review and editing

IGR: Conceptualization, Methodology, Validation, Analysis, Investigation, Data Curation, Visualization, Writing – original draft, Writing – review and editing

DLDC: Methodology, Validation, Investigation, Data Curation, Writing – review and editing

EKW: Methodology, Validation, Analysis, Data Curation, Visualization, Writing – original draft, Writing – review and editing

ELT: Methodology, Validation, Analysis, Data Curation, Visualization, Writing – original draft, Writing – review and editing

VAF: Conceptualization, Writing – original draft

JBR: Methodology, Validation, Investigation, Writing – review and editing KPW: Software, Writing – review and editing

JSS: Project Administration, Writing – review and editing

JLC: Conceptualization, Methodology, Project Administration, Resources, Supervision, Validation, Visualization, Writing – original draft, Writing – review and editing

## DATA AVAILABILITY

Databases can be found at https://doi.org/10.5281/zenodo.22019098. Raw counts for integrase activity are included in **Supplementary Table S5**.

## ACKNOWLEDGEMENTS

We thank the Cahill Laboratory members and the Sandia National Laboratories’ Environmental Systems Biology and Molecular and Microbiology Department for their valuable input during this study. The authors are grateful for the opportunity to demonstrate that this work could still succeed under conditions imposed by organizational changes. We thank Emily Hollister for reviewing a pre-submission manuscript draft, suggesting edits, and providing technical feedback. SandiaAI Chat, a version of OpenAI’s GPT-5.5 architecture, was used to assist with organization, drafting, and proofreading. All authors reviewed, approved, and take responsibility for the final manuscript.

