## Supplementary Data for "Temperature-Switchable Genome Editors from Extremophile-Derived Integrases"

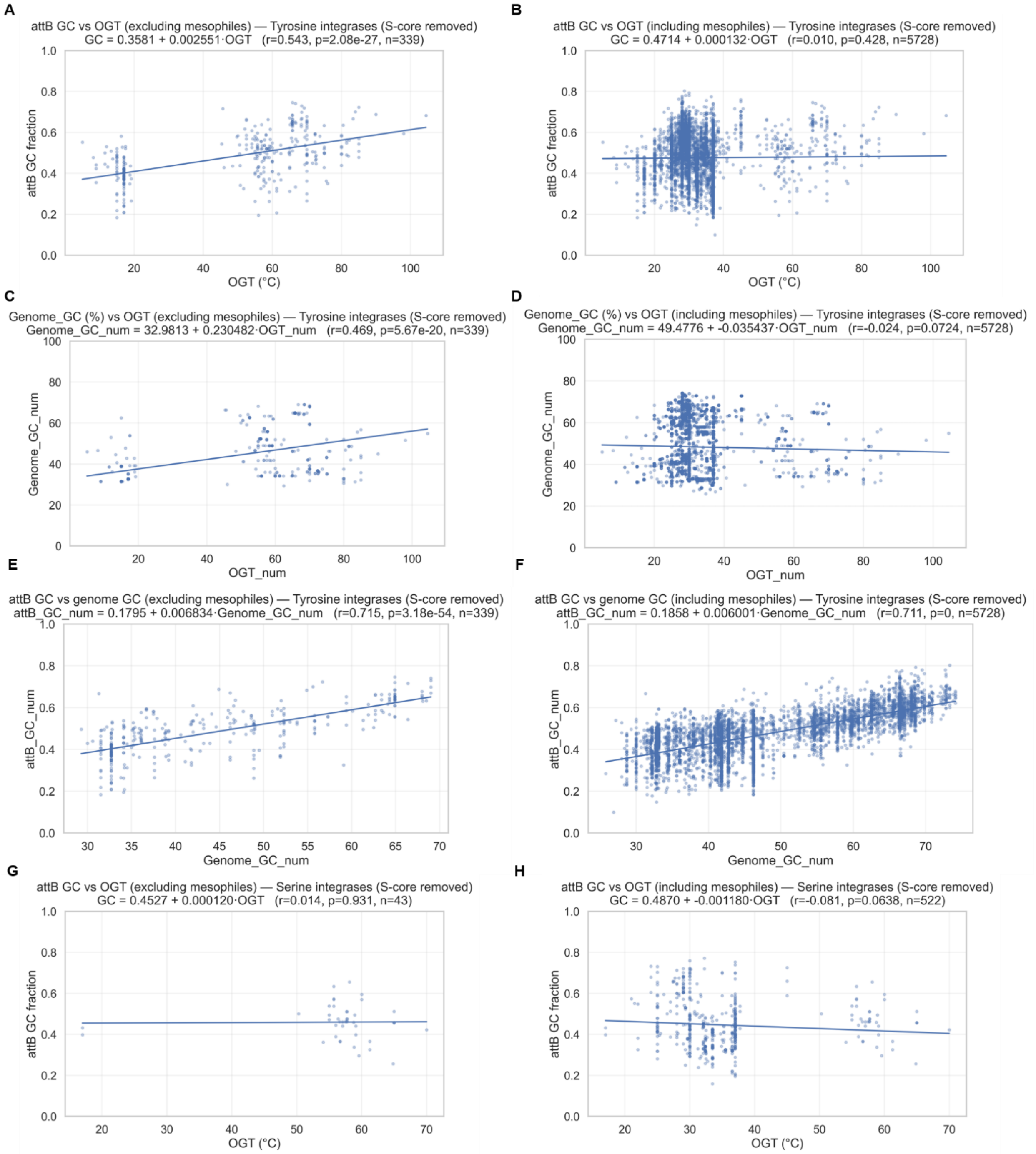

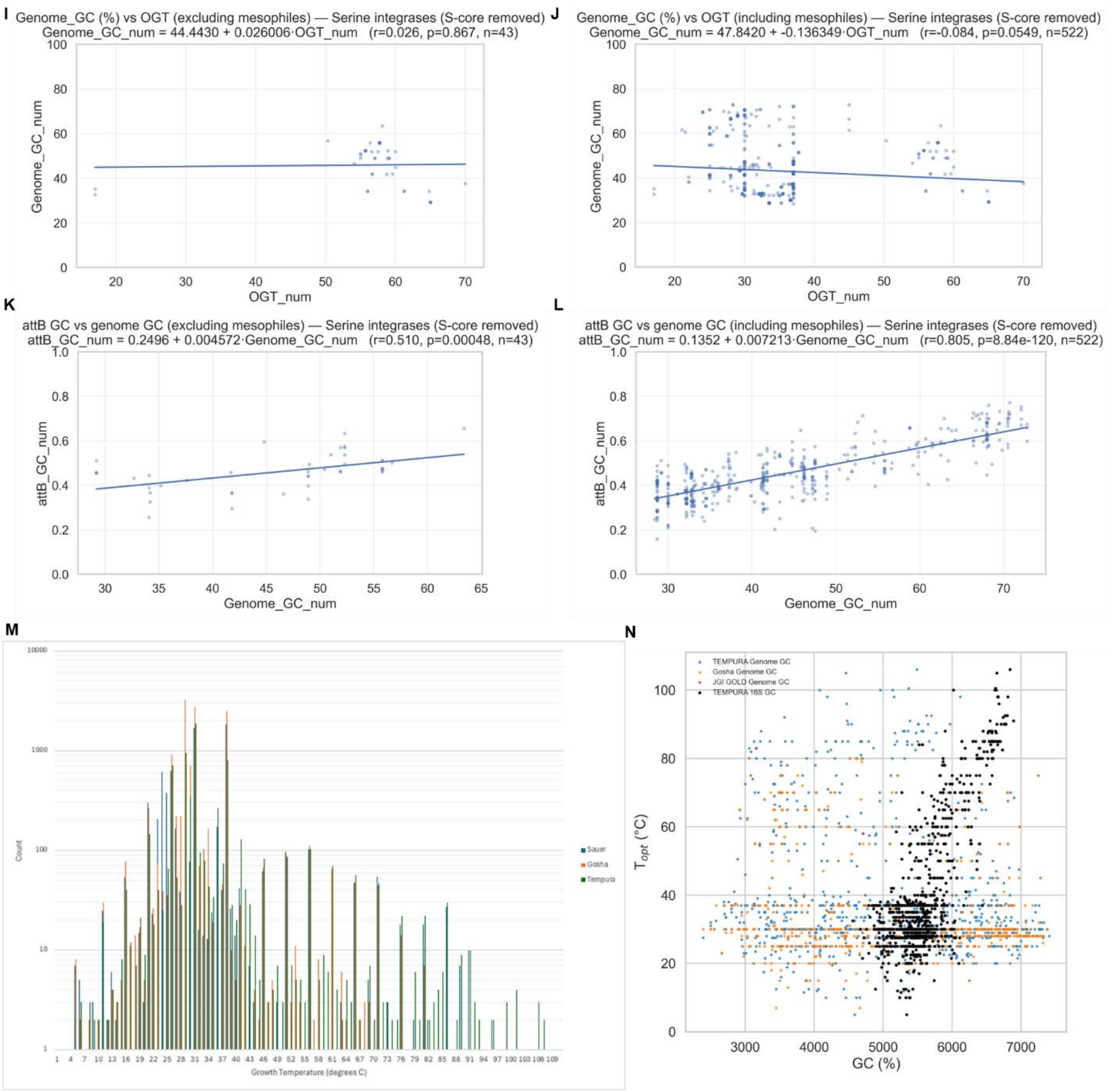


**Supplementary Figure S1. Characteristics of optimal growth temperature-linked database.** **(A-F)** Correlation plots for tyrosine integrase-associated GC regions. **(A-B)** *attB* GC vs OGT, **(C-D)** genome-wide GC vs OGT, and **(E-F)** *attB* GC vs genome-wide GC are shown excluding or including mesophile-class entries respectively (shown also in **Supp. Table S1**). **(G-L)** Correlation plots for serine integrase-associated GC regions. **(G-H)** *attB* GC vs OGT, **(I-J)** genome-wide GC vs OGT, and **(K-L)** *attB* GC vs genome-wide GC are shown excluding or including mesophile-class entries respectively (shown also in **Supp. Table S1**). **(M)** Count of species for which each temperature is reported as the optimal growth temperature (OGT). **(N)** A comparison of the OGT (˚C) and genomic GC content from TEMPURA (27), Gosha (26), and JGI GOLD (28) databases, indicating a lack of relationship between genomic GC content and OGT. In contrast, 16S gene GC content (black) from the TEMPURA database shows a positive relationship with increasing OGT. Linear regression analyses were performed in Python using the Scipy.stats package version 1.18.0 (30) with a significance threshold of $P<0.05$.

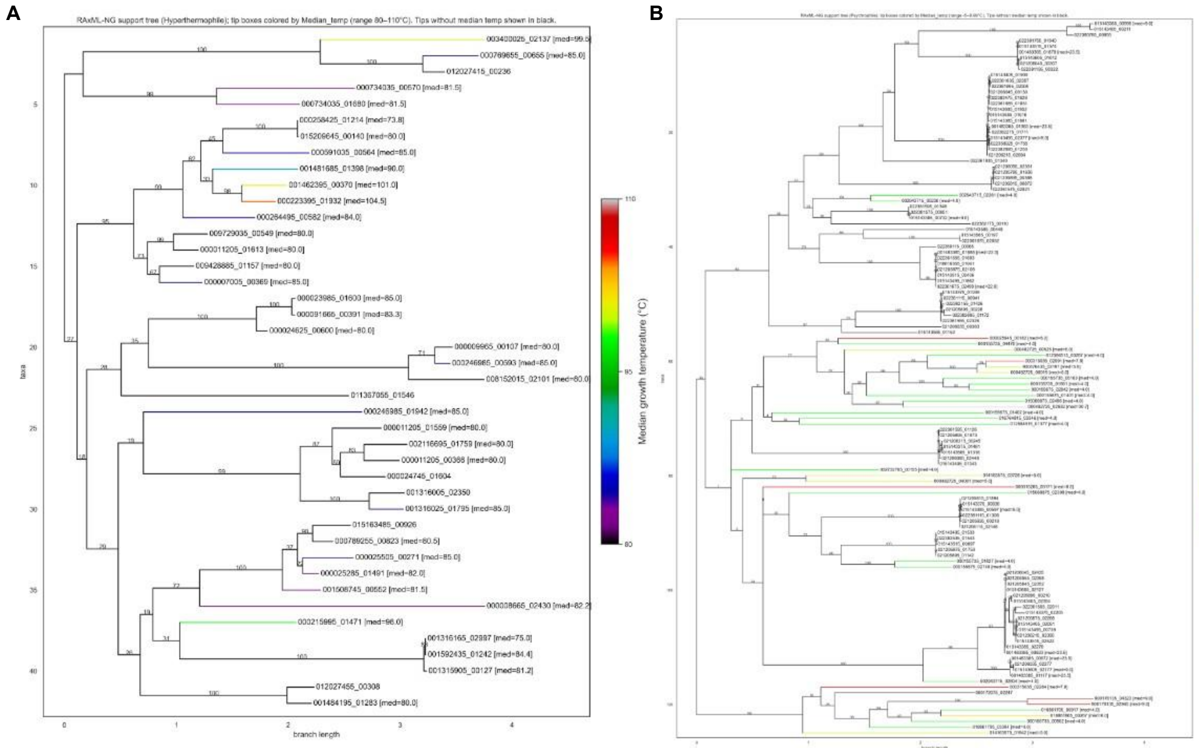


**Supplementary Figure S2. Examples of trees of thermally sorted integrases. (A)** Hyperthermophile (80-110˚C) and **(B)** psychrophile (-5-9.99˚C) trees were generated using RAxML-NG (33) to compare integrase similarity based on amino acid identity. Tip boxes are colored by median growth temperature.

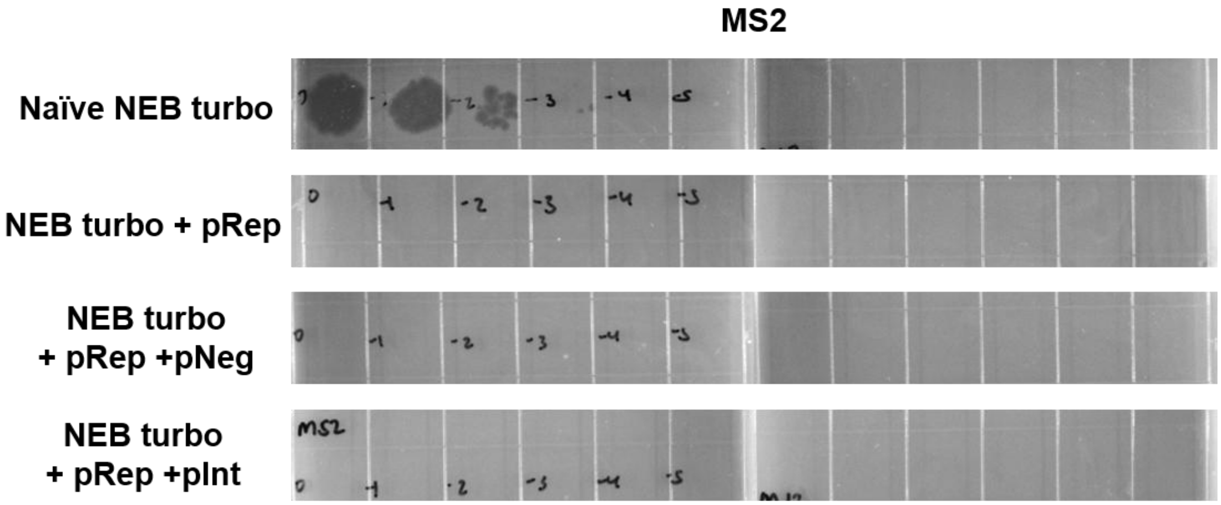


**Supplementary Figure S3. Plaque assays to evaluate F’ plasmid maintenance.** Plaque assays were performed with MS2 on naïve NEB turbo or NEB turbo containing candidate integrase plasmids. The native F’ plasmid in NEB turbo carries the *repE/incC* replication origin, as does pRep. The F pilus encoded by the native F’ plasmid is necessary for MS2 infection. Plaque assays are shown for all pRep variants.

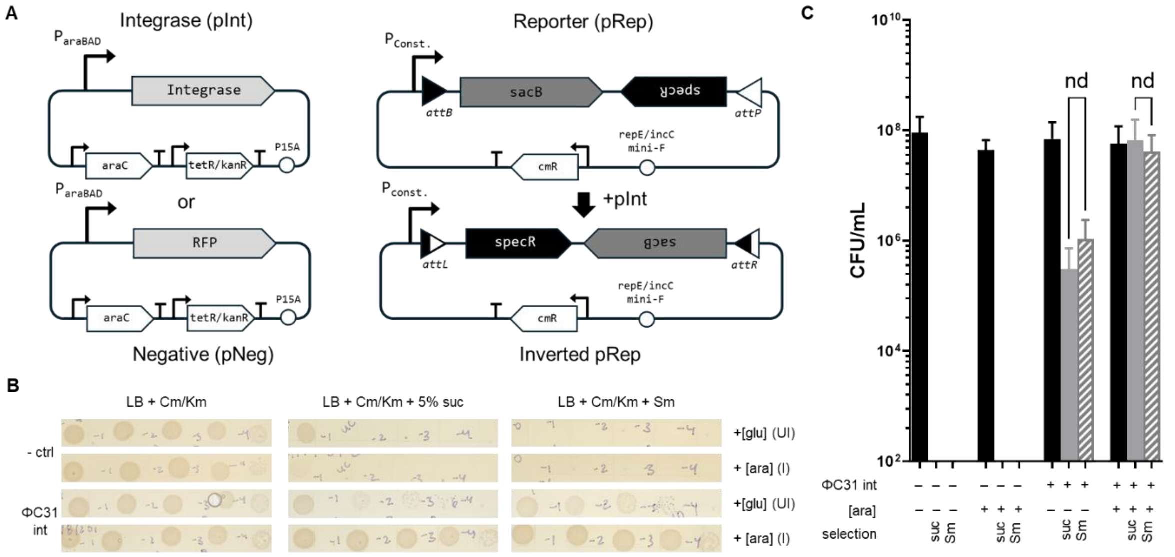


**Supplementary Figure S4. Integrase-mediated inversion testbed development.** A two-plasmid system **(A)** was used for assaying extremophile integrase activity. In the reporter plasmid, a strong constitutive promoter ($P_{16s}$) controls sacB-mediated counterselection pre-inversion and specR-mediated selection post-inversion event. This cassette is flanked by *attB* and *attP* sites which pair with a cognate integrase on a second plasmid under the control of an inducible promoter ($P_{araBAD}$). Serial dilutions were plated **(B)** and quantified **(C)** to assess integrase activity for the ΦC31 integrase and *att* pair. Integrase production was induced (arabinose, 0.2% w/v) or quenched (glucose, 0.2% w/v) and cells were plated on plates containing 5% w/v sucrose (solid grey bars) or 50 µg/mL spectinomycin (hatched grey bars) to select for inversion events or on plates that just contained selection for the two-plasmid system (solid black bars) (Cm/Tet). RFP was used as a negative control in place of the ΦC31 integrase to check for endogenous integrase-mediated inversion events. A representative example at 37˚C is shown and graph is normalized to limit of detection (100 CFU/mL). For statistical analysis, a Welch's t-test was performed in GraphPad Prism. *, P<0.05; **, P<0.01; ***, P<0.005; ****, P <0.0005.

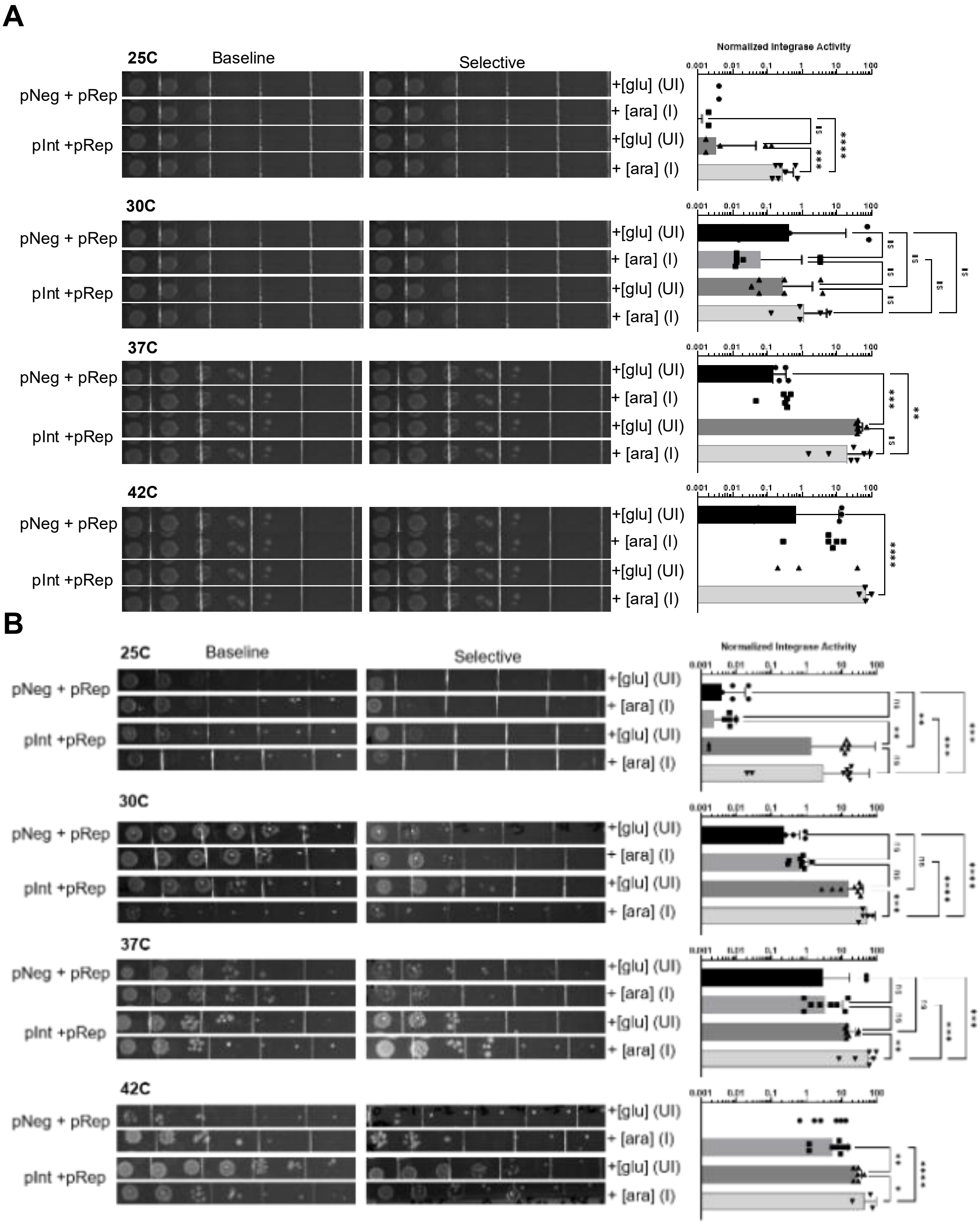


**Supplementary Figure S5. Inversion assays of extremophile *Geobacillus stearothermophilus*-derived integrases in *E. coli***. Serial dilutions were plated to assess integrase activity for the **(A)** *G. stearothermophilus* Gs1-derived, and **(B)** *G. stearothermophilus* Gs2-derived integrase and *att* pairs at four different temperatures (25˚C, 30˚C, 37˚C, and 42˚C). Integrase production was induced (arabinose, 0.2% w/v) or quenched (glucose, 0.2% w/v) and cells were plated on plates containing 5% w/v sucrose to select for inversion events or on plates that just contained selection for the two-plasmid system (Cm/Tet). RFP was used as a negative control in place of integrases to check for endogenous integrase-mediated inversion events. For statistical analysis an ordinary one-way ANOVA was performed in GraphPad Prism. *, $P<0.05$; **, $P<0.01$; ***, $P<0.001$; ****, $P <0.0001$.

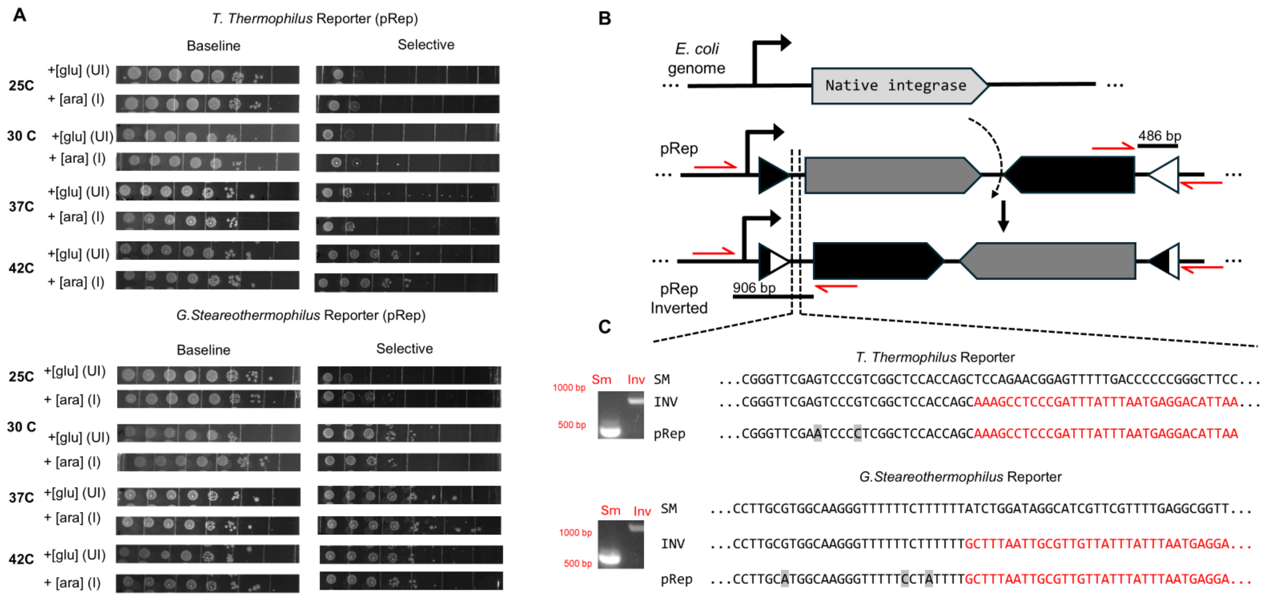


**Supplemental Figure S6. Endogenous *E. coli* integrases can act on extremophile *att* sites. (A)** Transformants containing only reporter plasmids (pRep) were plated at four temperatures (25˚C, 30˚C, 37˚C, and 42˚C) on baseline (Cm/Tet) and selective (Cm/Tet 5% w/v Sucrose) conditions to determine endogenous activity on *T. thermophilus*-derived or *G. stearothermophilus*-derived *att* sites. **(B)** Proposed mechanism of a native *E. coli* integrase catalyzing the inversion of pRep between attachment sites. Molecular confirmation of inversion events *via* colony PCR using primer pairs (**Supp. Table S3**) that faced outward on the inversion cassette and inward on the plasmid backbone. **(C)** The expected band size for starting material was approximately 486 bp and approximately 906 bp for inversion events. PCR and agarose gel electrophoresis showed bands at the expected size for starting material (SM) and inversion (Inv) both reporters. Nanopore sequencing confirmed that pRep alone for both reporters aligned with the inverted sequence (red text in sequence) that would be expected upon integrase-mediated inversion activity.
